# Comparison of urine proteomes from tumor-bearing mice with those from tumor-resected mice

**DOI:** 10.1101/2022.06.08.495253

**Authors:** Heng Ziqi, Zhao Chenyang, Gao Youhe

## Abstract

**[Objective]:** This study focuses on the most important concern of surgeons - whether they resected all of the tumor. Urine can reflect early changes associated with physiological or pathophysiological processes. Based on the above ideas, we conducted experiments to explore changes in the urine proteome between tumor-bearing mice and tumor-resected mice.

**[Method]:** The tumor-bearing mouse model was established with MC38 mouse colon cancer cells, and the mice were divided into the healthy control group, complete resection group, and nonresection group. Urine was collected 7 days and 30 days after resection. Liquid chromatography coupled with tandem mass spectrometry (LC–MS/MS) was used to identify the urine proteome and then analyze differentially expressed proteins and biological pathways.

**[Results]:** (1) Seven days after tumor resection, there were 20 differentially expressed proteins that could distinguish between the complete resection group and the nonresection group. The biological process includes circadian rhythm, Notch signaling pathway, leukocyte cell–cell adhesion, and heterophilic cell–cell adhesion via plasma membrane cell adhesion molecules. (2) Thirty days after tumor resection, there were 33 differentially expressed proteins that could distinguish between the complete resection group and the nonresection group. The biological process includes cell adhesion, complement activation, the alternative pathway, the immune system process, and angiogenesis. (3) There was no significant difference between the two groups at 30 days after tumor resection between the complete resection group and the healthy control group.

**[Conclusion]:** Changes in the urine proteome can reflect tumors with or without complete resection.

## 1 Introduction

The most common method of surgical treatment for solid tumors is resection with chemotherapy or radiotherapy, but recurrence of tumors after treatment is very common. Whether the tumor is resected cleanly is a major concern for many surgeons.

Every cell in the body depends on a stable internal environment to survive and function. Blood, as the provider of the internal environment, needs to be stable and balanced to protect cells from disturbing factors. In contrast, as a filtrate of blood, urine does not need or have a stabilizing mechanism and is not regulated by homeostatic mechanisms. It is able to enrich for changes caused by the body’s disease in its early stage. Thus, urine is a good biological source for finding biomarkers^[1]^.

In previous studies, the urinary proteome of Walker 256 tumor-bearing rats showed significant changes prior to the growth of palpable tumor masses, and these early changes in the urine could also be identified by differential abundance in the late stages of cancer^[2]^. Changes in the urinary proteome occurred on Day 2 after tail vein injection of Walkers-256 cells in rats, earlier than the pathological changes in lung tumor nodules appeared on Day 4^[3]^. Twenty-five proteins in the urine of rats in the tumor group changed significantly 3 days after Walker 256 cells were implanted in the tibia of rats, predating the detection of significant lesions on computed tomography (CT) scans^[4]^. On Day 3 after liver injection using Walker-256 cells, 12 proteins were significantly changed in the experimental rats, 7 proteins were significantly associated with liver cancer, and the same tumor cells growing in different organs were reflected in the differential urine proteins^[5]^.

There are many factors affecting urine samples, and it is time-consuming to collect samples from early-stage patients, so establishing a mouse model of colorectal cancer minimizes potential interfering factors and allows dynamic monitoring of disease progression and obtaining urine samples prior to pathology or clinical presentation, facilitating the observation of tumor resection and recurrence ^[6]^.

In this study, MC38 cell line derived from C57BL6 murine colon adenocarcinoma subcutaneous tumor model was established. Then, the tumors were resected from the tumor-bearing mice. Urine samples were collected after the surgery, and liquid chromatography-tandem mass spectrometry (LC– MS/MS) was performed for urine proteomics analysis. The workflow is shown in Figure 1. The aim of this study was to compare urine proteomes from tumor-bearing mice with those from tumor-resected mice.

**Figure 1.**
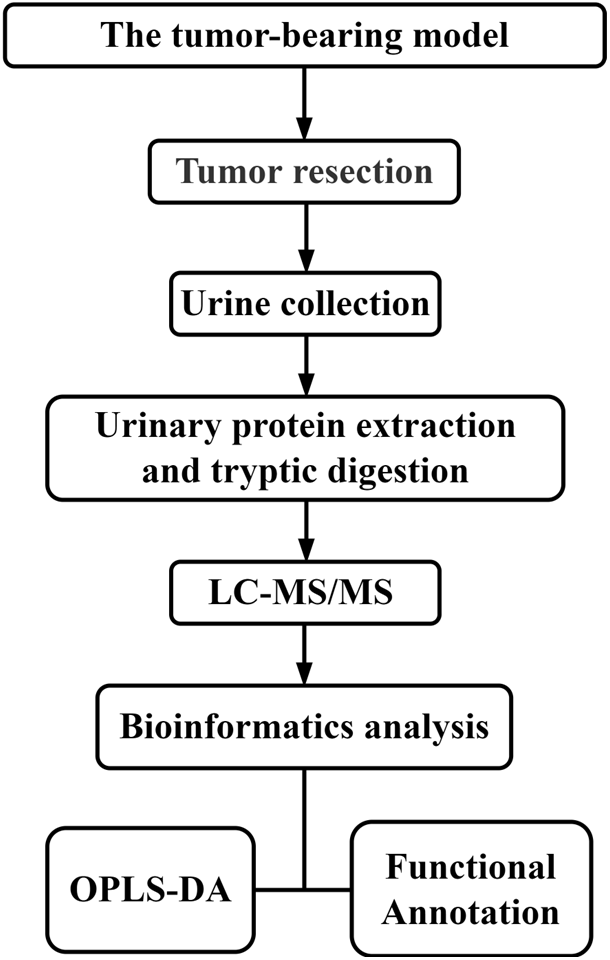
The experimental design and workflow of the analysis in this study The tumor-bearing model was established, and then the tumors were resected from the tumor-bearing mice. Urine samples were collected at Days 7 and 30 after tumor resection. Urine proteins were identified by liquid chromatography coupled with tandem mass spectrometry (LC–MS/MS).

## 2 Materials and Methods

### 2.1 Establishment of the MC38 tumor-bearing mouse model

Male c57BL/6j mice (18 g-20 g) were supplied by Beijing Vital River Laboratory Animal Technology Co., Ltd. All animals were maintained with free access to a standard laboratory diet and water with a 12-h light–dark cycle under controlled indoor temperature (22 ± 2 °C) and humidity (65– 70%). All methods in this research were performed in accordance with the guidelines and regulations. The experimental mice were kept in the new environment for three days to familiarize them with the environment. Then, they were randomly divided into three groups: a healthy control group (n=5), a complete resection group (n=5) and a nonresection group (n=5). MC38 cells (provided by Prof. Sheng’s research group at Beijing University of Technology) were added to complete culture medium (DMEM, 10% fetal bovine serum and 1% penicillin/streptomycin) and placed in T75 cell culture flasks. After sufficient cells were prepared, the MC38 tumor cells were collected, centrifuged, and resuspended in phosphate-buffered saline (PBS) for the subsequent establishment of the mouse models. Cell viability was assessed by the trypan blue exclusion test. MC38 cells were stained with 0.4% trypan blue solution and then counted using a hemocytometer. It was observed that more than 95% of the tumor cells were viable.

The mice were anesthetized with sodium pentobarbital at a dose of 4 mg/kg. After they were anesthetized, the mice were fixed on disposable sterile medical pads, and the injection site was disinfected after removing the hair. The experimental mice in the complete resection group (n=5) and nonresection group (n=5) were subcutaneously inoculated with 5 × 10^6^ viable MC38 cells in 200 μl of PBS into the right hind limb of the animal. The healthy control group (n=5) was injected with PBS buffer into the same position. The mice in each of the three groups underwent resection after establishment of the tumors. The subcutaneous tumor was completely resected from the mice in the complete resection group; the subcutaneous tumor was preserved in the nonresection group, and only part of the tissue and muscle were removed to ensure the same wound surface as the mice in the other two groups; only part of the tissue and muscle were removed in the healthy control group to ensure the same wound surface as the mice in the other two groups.

### 2.2 Urine sample collection and preparation

Urine was collected from mice in the three groups on Days 7 and 30 after resection. During urine collection, all mice were placed individually in metabolic cages with no food or water. Urine was collected overnight for 12 hours, and the volume of urine collected was not less than 1 ml. The urine samples were stored at –80 °C immediately after collection.

Urine samples were defrosted at 4 °C and centrifuged at 12,000 × g for 30 min at 4 °C to remove cell debris. Then, the supernatants were precipitated with three volumes of ethanol at -20 °C, followed by centrifugation at 12,000 × g for 30 min. The pellet was resuspended in lysis buffer (8 mol/L urea, 2 mol/L thiourea, 50 mmol/L Tris, and 25 mmol/L DTT). The protein concentration of each sample was measured using the Bradford assay. The protein samples were stored at −80 °C for later use.

The urinary proteins were prepared using the FASP method^[7]^, and then peptides were collected after enzymatic digestion using trypsin (Trypsin Gold, Promega, Fitchburg, WI, USA); the peptides were desalted by Oasis HLB cartridges (Waters, Milford, MA) and then drained using a vacuum drier. The peptides were redissolved in 0.1% formic acid water and diluted to 0.5 μg/μL. A mixed peptide sample was prepared from each sample, graded using a high pH reversed-phase peptide separation kit (Thermo Fisher Scientific), evacuated using a vacuum drier and then redissolved in 0.1% formic acid in water for subsequent library construction. The iRT (Biognosys) was added to all identified samples for retention time uniformity.

### 2.3 LC–MS/MS analysis

The peptides were separated by using an EASY-nLC 1200 UPLC (Thermo Fisher Scientific, USA) system. Peptides were dissolved in 0.1% formic acid in water, and then 1 μg of each peptide sample was loaded on a rap column (75 µm × 2 cm, 3 µm, C18, Thermo Fisher). The eluate was loaded onto a reversed-phase analytical column (50 µm × 250 mm, 2 µm, C18, Thermo Fisher) with an elution gradient of 4%-35% mobile phase B (80% acetonitrile + 0.1% formic acid + 20% water at a flow rate of 300 nL/min) for 90 min. For fully automated and sensitive signal processing, a calibration kit (iRT kit, Biognosys, Switzerland) was used with all samples at a concentration of 1:20 v/v. Then, fractions were analyzed in DDA-MS mode with the following settings: 2.4 kV for spray voltage, 60000 for Orbitrap primary resolution, 350-1550 m/z for scan range, 200-2000 m/z for secondary scan range, 30000 for resolution, 2 Da for screening window, 30% HCD for collision energy. The AGC target was 5e4, and the maximum injection time was 30 ms. The raw files were used to build a dataset and were analyzed by PD software (Proteome Discoverer 2.1, Thermo Fisher Scientific, Inc.).

The PD search results were used to establish the DIA acquisition method, and the m/z distribution density was used to calculate the window width and number. A single peptide sample was subjected to DIA mode mass spectrometry acquisition, and each sample was repeated twice. Thirty samples were analyzed by DIA-MS mode. The liquid phase parameters were acquired in the same way as in the DDA database. The parameters of the mass spectrometry were set as follows: first level full scan at 60,000 resolution, 350-1550 m/z for scan range, followed by a second level scan at 30,000 resolution, 39 screening windows, 30% HCD for collision energy, AGC target of 1e^6^, and 50 ms for maximum injection time. Window calculation: the DDA search results from the library acquisition were sorted into 39 groups based on m/z. The m/z range of each group is the window width of the collected DIA data. During the sample analysis, a mixture from each sample was analyzed after every seven samples for quality control (QC).

### 2.4 Mass spectrometry data analysis

MS data were processed and analyzed using Spectronaut software. The raw files acquired by DIA for each sample were imported for the library search. A highly credible protein standard was peptide *q* < 0.01, and all fragment ion peak areas of secondary peptides were used for protein quantification.

### 2.5 Statistical analysis

The k-nearest neighbor (K-NN) method was used to fill the missing values of protein abundance. Comparisons between two groups were performed by one-way ANOVA. The differential proteins at Days 7 and 30 were screened by the following criteria: fold change ≥ 1.5 or fold change ≤ 0.67. Group differences resulting in *p* <0.05 were identified as statistically significant.

### 2.6 Randomized grouping statistical analysis

To determine that the differential proteins identified were due to random allocation, the same criteria for screening differential proteins were applied: fold change ≥ 1.5 or fold change ≤ 0.67 and *p* <0.05.

### 2.7 Functional analysis of differentially expressed proteins

We used the ‘Wu Kong’ platform (https://www.omicsolution.com/wkomics/main/) relative orthogonal signal-corrected partial least squares discriminant analysis (OPLS-DA) ^[8]^. The differential proteins at the different time points were analyzed by using the DAVID database (https://david.ncifcrf.gov/) to determine the functional annotation (*p* < 0.05 was identified as statistically significant).

## 3 Results and Discussion

### 3.1 Establishment of animal models

All three groups underwent resection 7 or 8 days after tumor cell inoculation, when the tumors had grown to a palpable state (the method is shown above), and urine was collected on Days 7 and 30 after resection. The procedure is shown in Figure 2.

**Figure 2.**
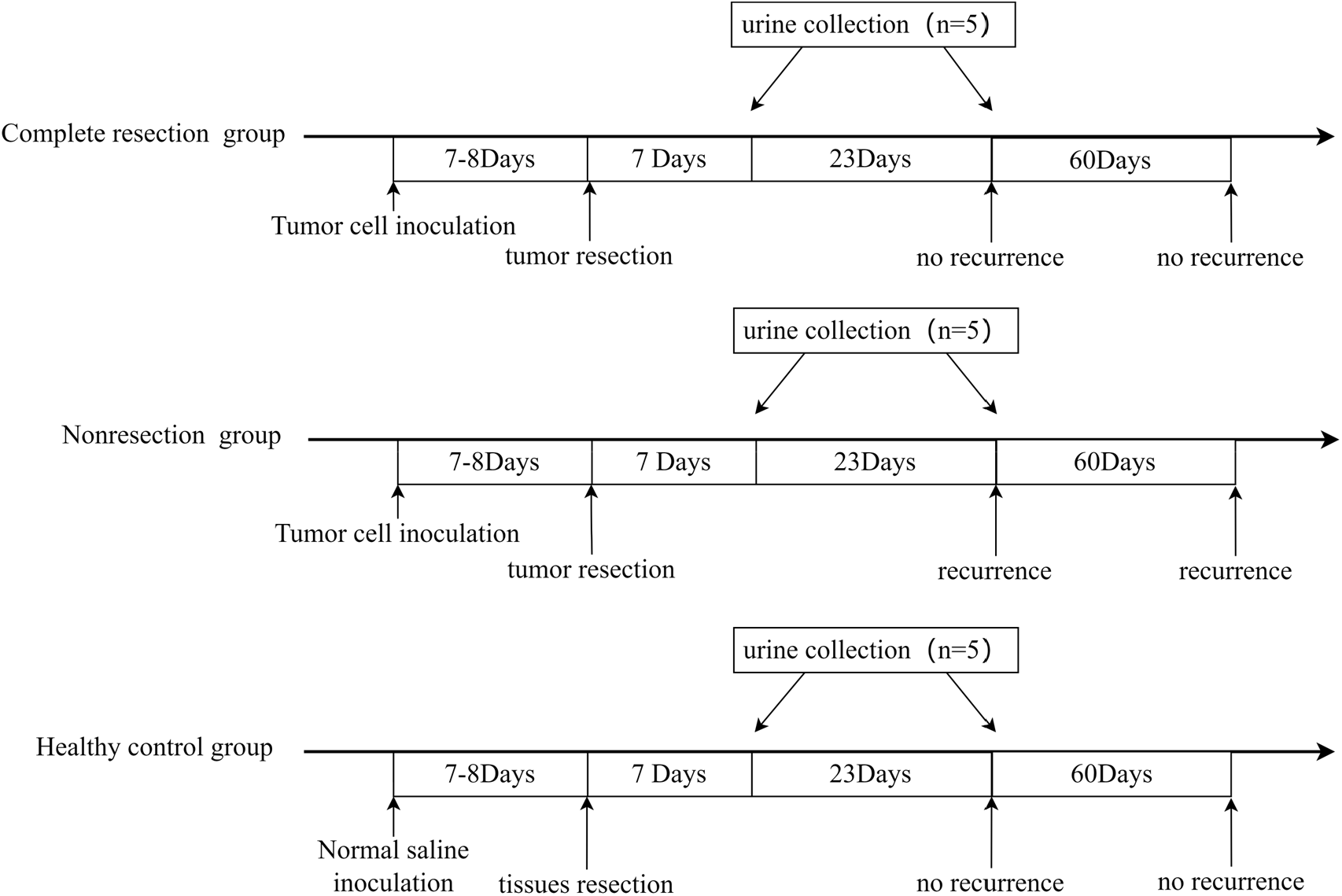
Animal model establishment process and the time point of urine collection

Animal models were successfully established, and the tumor size at 7 or 8 days was as shown in Table 1. No recurrence was seen 90 days after resection in the complete resection group. The resection was considered successful. No abnormalities were seen in the healthy control group.

**Table 1.** Tumor size (expressed as the mean diameter)

| Complete resection group (n=5) |  |  |  |  |  |
| --- | --- | --- | --- | --- | --- |
| No. | 1 | 2 | 3 | 4 | 5 |
| Size (mm) | 4.2 | 5.0 | 5.2 | 4.6 | 4.5 |
| Nonresection group (n=5) |  |  |  |  |  |
| No. | 6 | 7 | 8 | 9 | 10 |
| Size (mm) | 4.6 | 3.7 | 6.0 | 4.8 | 3.4 |
| Healthy control group (n=5) |  |  |  |  |  |
| No. | 11 | 12 | 13 | 14 | 15 |
| size(mm) | - | - | - | - | - |

### 3.2 Analysis of urine proteome changes

#### 3.2.1 Orthogonal partial least-squares discrimination analysis (OPLS-DA) for total urine protein at different time points

Analysis of the results of the urine proteome at two time points for the three groups showed that a total of 405 proteins were identified in all samples.

Using OPLS-DA, the total urinary proteins identified in the complete resection group and nonresection group (n=10) at Day 7 were analyzed, and the results are shown in Figure 3(a). The two groups can be clearly distinguished. The values of R^X, R^Y and Q^Y were 0.369, 0.993 and 0.788, respectively, and the data indicated good model fit accuracy. The Variable Importance for the Projection (VIP) was calculated for the differential proteins, and it was found that all 20 differential proteins met the criteria of VIP value > 1.0. The results indicate that the differential proteins have a strong ability to discriminate between the two groups.

**Figure 3.**
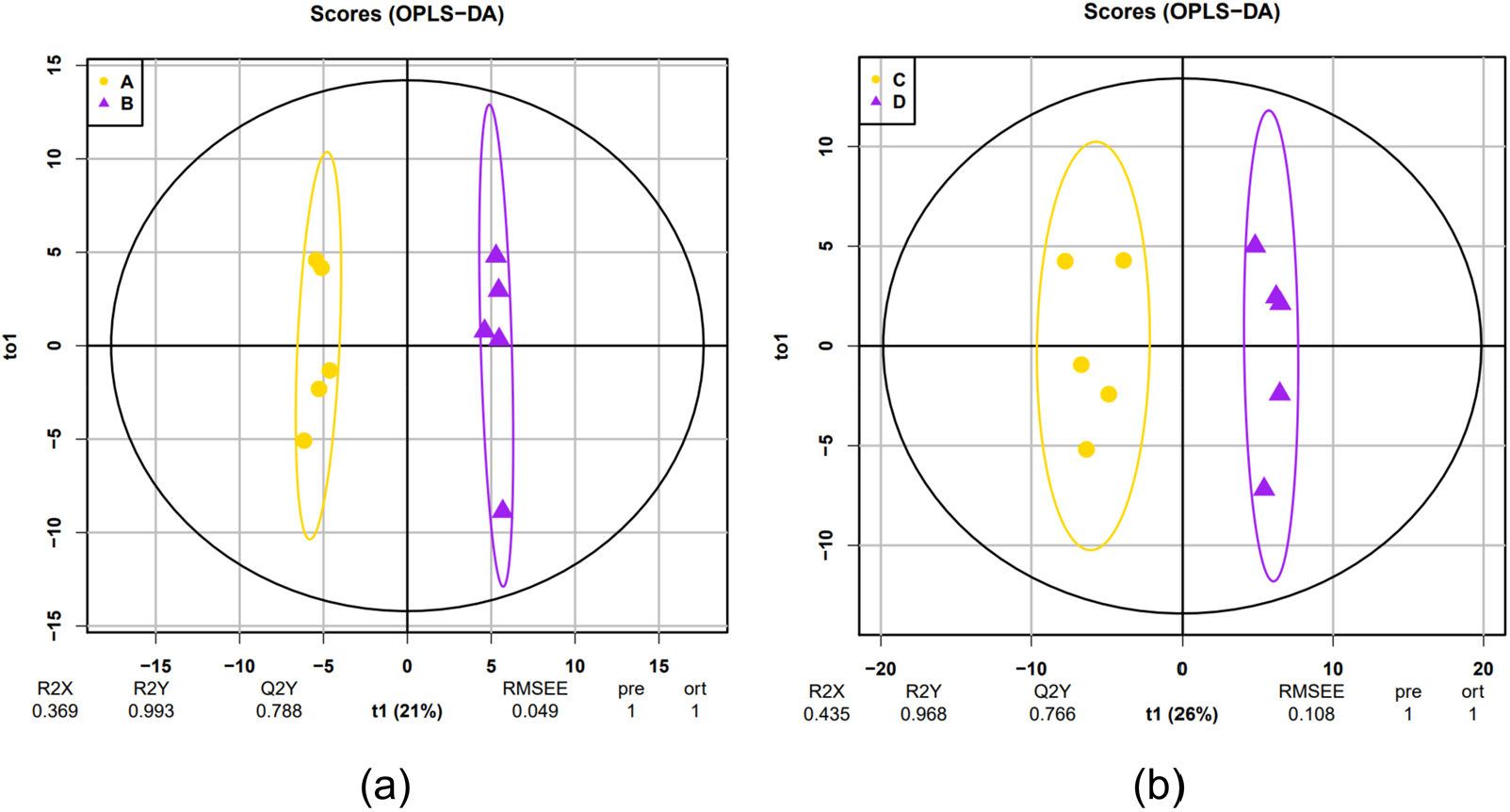
The results of OPLS-DA A: complete resection group-D7; B: nonresection group-D7; C: complete resection group-D30; D: nonresection group-D30

Similarly, the total urinary proteins identified in the complete resection group and nonresection group (n=10) at Day 30 were analyzed by OPLS-DA, and the results are shown in Figure 3(b). The two groups can be clearly distinguished. The values of R^X, R^Y and Q^Y were 0.435, 0.968 and 0.766, respectively, and the model fit accuracy was good. The VIP calculation for the difference proteins revealed that all 33 difference proteins met the criteria of VIP value > 1.0. The results indicate that the differential proteins had a strong ability to discriminate between the two groups.

#### 3.2.2 Differential protein and biological pathways on Day 7 after resection between the complete resection group and the nonresection group

There were 20 differential proteins (shown in Table 2) identified with statistically significant changes between the complete resection group and the nonresection group at Day 7 after resection, of which 9 differential proteins were upregulated and 11 differential proteins were downregulated. Considering that omics data are large but the sample size is limited, the differences between the two groups may be randomly generated. To confirm whether the differential proteins were indeed due to resection, we randomly allocated the proteomic data of 10 samples at each time point. Random grouping statistical analysis showed that an average of approximately 3.44 differential proteins were identified out of 125 random grouping results, indicating that at least 83% of the differential proteins were due to resection of the subcutaneous tumor.

**Table 2.**
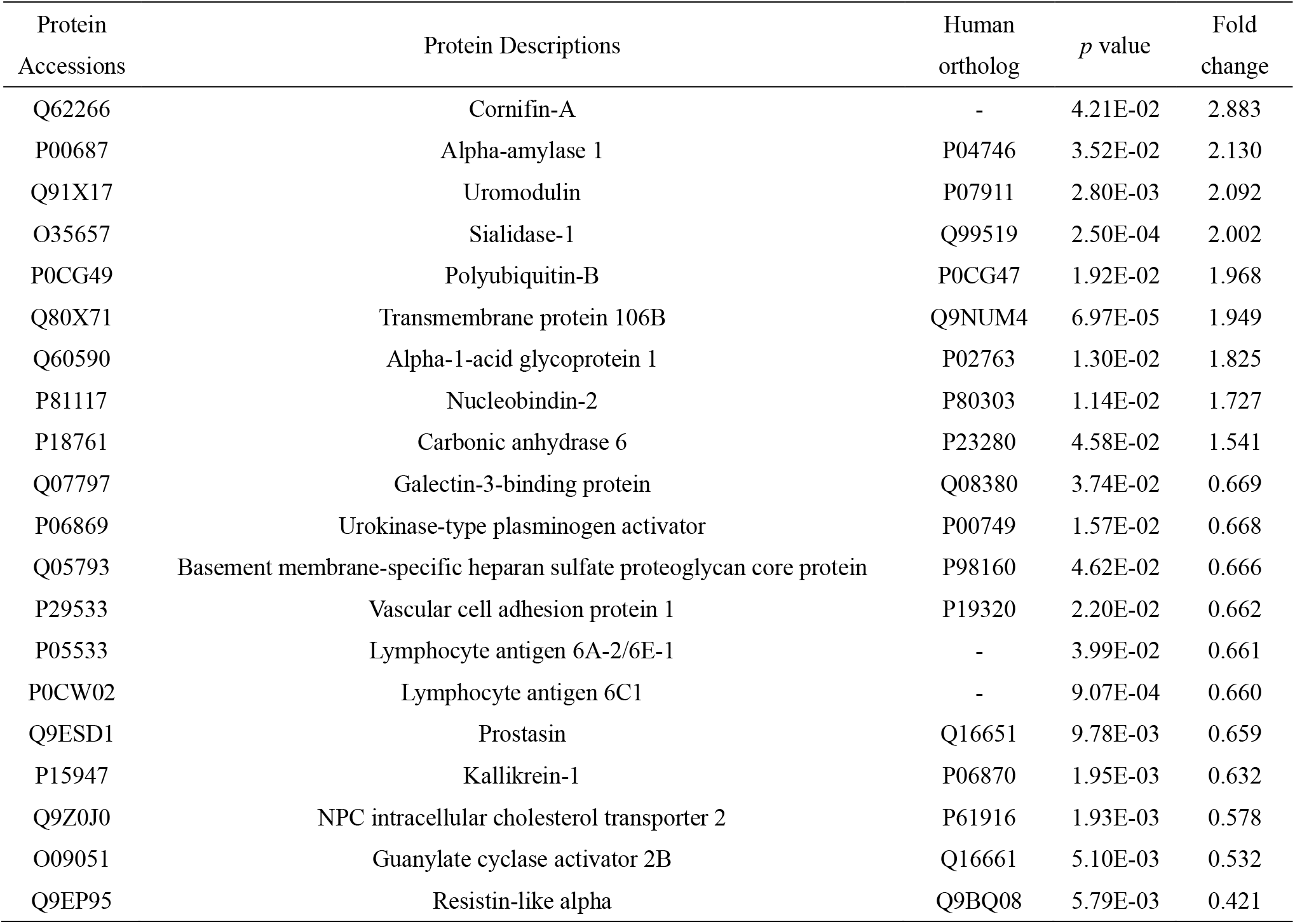
Differential proteins on Day 7 between the complete resection group and the nonresection group

Six biological processes were enriched by differential proteins (shown in Figure 4), including circadian rhythm, Notch signaling pathway, leukocyte cell–cell adhesion, and heterophilic cell–cell adhesion via plasma membrane cell adhesion molecules. Some studies have shown that circadian rhythms are relevant to both wound healing and tumor growth, such as wound healing accompanied by inflammation, leukocyte transport, and tissue remodeling, and that some of these proteins are involved in a circadian-driven chronological coordination mechanism^[9, 10]^. The Notch signaling pathway is involved in the regulation of multidimensional subcutaneous tumorigenic behavior^[11]^. Notch signaling is associated with oncogene expression, has the potential to be a target for tumor suppressors and affects tumor cell proliferation, differentiation, apoptosis and genomic instability^[12]^. Regulation of lymphatic endothelial cell expression of the integrin ligands ICAM-1 and VCAM, which control leukocyte transition, induces enhanced leukocyte adhesion to generate an immune response against tumors^[13]^.

**Figure 4.**
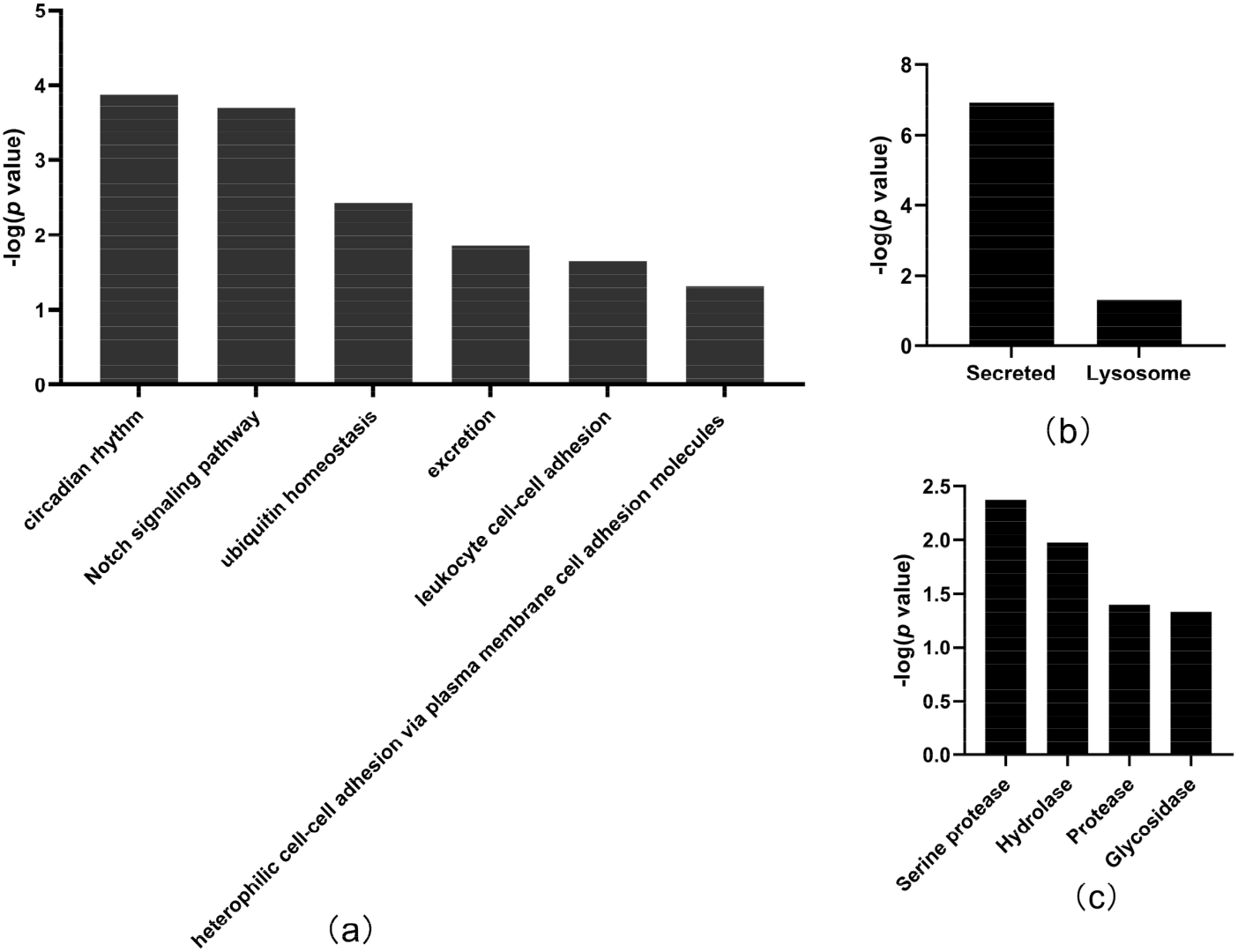
Biological process (a), cellular component (b) and molecular function (c) in Day 7 between the complete resection group and the nonresection group

In the cellular component category (Figure 4 (b)), all of these differential proteins were from secreted and lysosome. In the molecular function category (Figure 4 (c)), serine protease and hydrolase were overrepresented.

#### 3.2.3 Differential protein and biological pathways on Day 30 after resection between the complete resection group and the nonresection group

Thirty-three differential proteins were identified between the complete resection group and the nonresection group at Day 30 after surgery (as shown in Table 3), and 13 proteins were upregulated and 20 proteins were downregulated, enriched 19 biological processes (as shown in Figure 5). Random grouping statistical analysis showed that an average of approximately 3.76 differential proteins were identified out of 125 random grouping results, indicating that at least 89% of the differential proteins were due to differences in the level of resection of the tumor.

**Table 3.** Differential proteins on Day 30 between the complete resection group and the nonresection group

| Protein<br>Accessions | Protein Descriptions | Human<br>ortholog | *p* value | Fold<br>change |
| --- | --- | --- | --- | --- |
| Q8VED5 | Keratin, type II cytoskeletal 79 | Q5XKE5 | 2.81E-02 | 3.026 |
| P00687 | Alpha-amylase 1 | P04746 | 2.16E-02 | 2.778 |
| Q922U2 | Keratin, type II cytoskeletal 5 | P13647 | 1.10E-02 | 2.732 |
| Q61781 | Keratin, type I cytoskeletal 14 | P02533 | 2.12E-02 | 2.685 |
| P11591 | Major urinary protein 5 | - | 2.94E-02 | 2.292 |
| Q6NXH9 | Keratin, type II cytoskeletal 73 | Q86Y46 | 4.80E-02 | 1.997 |
| P03953 | Complement Factor D | P00746 | 8.82E-04 | 1.784 |
| O55186 | CD59A glycoprotein | P13987 | 2.11E-02 | 1.671 |
| Q61581 | Insulin-like growth factor-binding protein 7 | Q16270 | 2.78E-03 | 1.598 |
| P09803 | Cadherin-1 | P12830 | 5.21E-03 | 1.542 |
| Q9JJS0 | Signal peptide, CUB and EGF-like domain-containing protein 2 | Q9NQ36 | 5.10E-03 | 1.535 |
| P11589 | Major urinary protein 2 | - | 3.36E-02 | 1.523 |
| P01132 | Pro-epidermal growth factor | P01133 | 3.61E-03 | 1.516 |
| Q07456 | Protein AMBP | P02760 | 2.47E-03 | 0.652 |
| P47878 | Insulin-like growth factor-binding protein 3 | P17936 | 8.47E-03 | 0.628 |
| P04186 | Complement factor B | P00751 | 4.87E-02 | 0.627 |
| Q9WTR5 | Cadherin-13 | P55290 | 3.56E-02 | 0.617 |
| P11276 | Fibronectin | P02751 | 2.30E-03 | 0.594 |
| Q61129 | Complement factor I | P05156 | 1.51E-02 | 0.579 |
| Q921W8 | Secreted and transmembrane protein 1A | Q8WVN6 | 3.60E-04 | 0.575 |
| O88968 | Transcobalamin-2 | P20062 | 2.67E-02 | 0.538 |
| Q91X72 | Hemopexin | P02790 | 1.94E-03 | 0.473 |
| P0CW02 | Lymphocyte antigen 6C1 | - | 1.21E-03 | 0.466 |
| Q8K4G1 | Latent-transforming growth factor beta-binding protein 4 | Q8N2S1 | 1.15E-02 | 0.454 |
| O09051 | Guanylate cyclase activator 2B | Q16661 | 2.69E-03 | 0.442 |
| P25119 | Tumor necrosis factor receptor superfamily member 1B | P20333 | 2.37E-02 | 0.431 |
| Q9EP95 | Resistin-like alpha | Q9BQ08 | 9.20E-03 | 0.389 |
| Q05793 | Basement membrane-specific heparan sulfate proteoglycan core protein | P98160 | 4.91E-04 | 0.378 |
| P29533 | Vascular cell adhesion protein 1 | P19320 | 1.16E-02 | 0.352 |
| O08997 | Copper transport protein ATOX1 | O00244 | 1.02E-02 | 0.318 |
| Q91VW3 | SH3 domain-binding glutamic acid-rich-like protein 3 | Q9H299 | 3.02E-02 | 0.297 |
| Q4KML4 | Costars family protein ABRACL | Q9P1F3 | 3.05E-02 | 0.254 |
| Q61646 | Haptoglobin | P00739 | 2.19E-02 | 0.214 |

**Figure 5.**
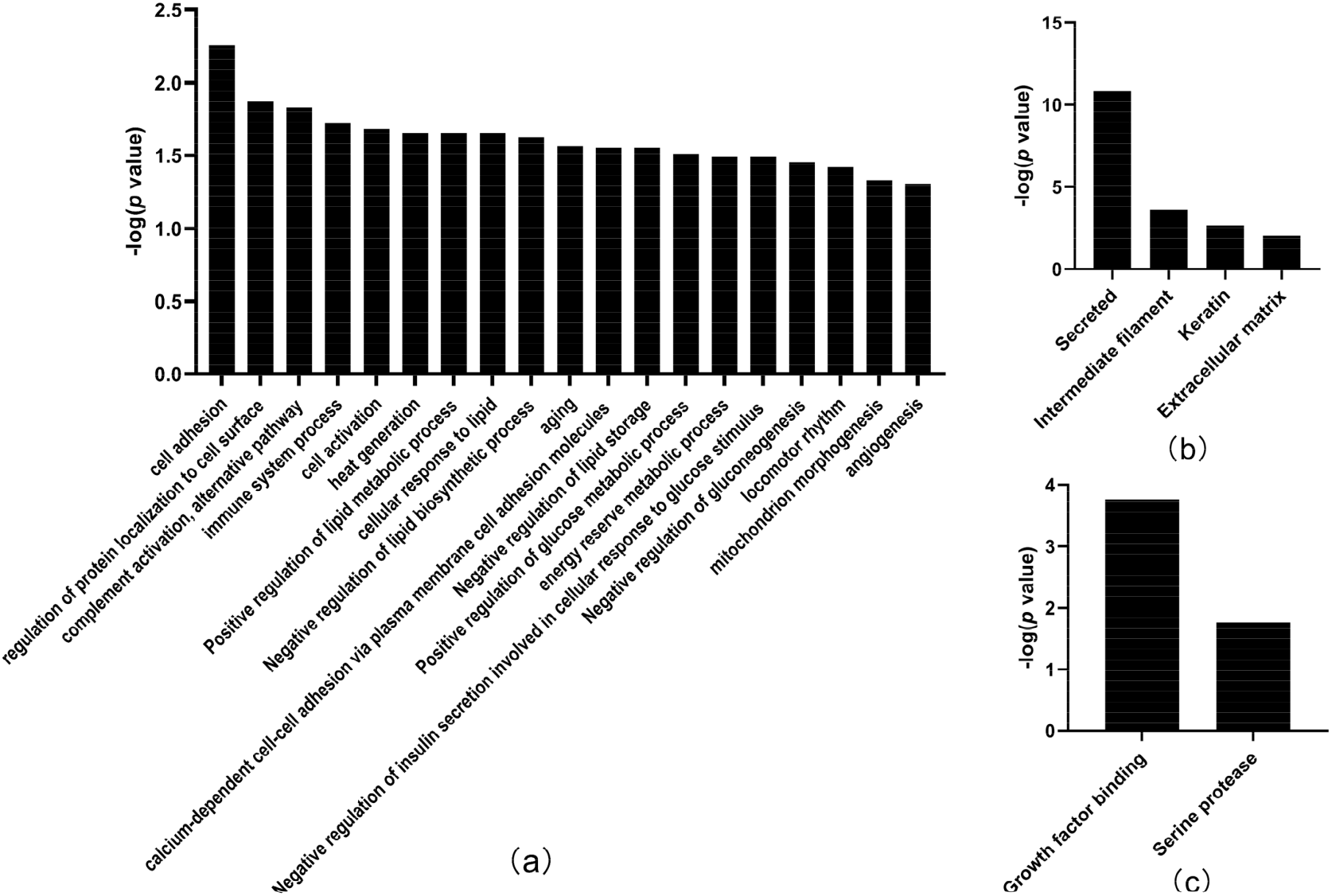
Biological process (a), cellular component and molecular function (c) on Day 30 between the complete resection group and the nonresection group

The biological processes included cell adhesion, regulation of protein localization to the cell surface, complement activation, alternative pathways, immune system processes, positive regulation of glucose metabolic processes, mitochondrion morphogenesis, and angiogenesis. Among them, it has been reported that cell adhesion and regulation of protein localization to the cell surface are relevant to tumor growth, especially cadherin and integrin, as these proteins act as receptors for ligand activation and activate relevant signaling through changes in the physical environment^[14]^. Immune-related pathways are frequently altered during tumor growth, such as complement activation in the tumor microenvironment, which enhances tumor growth and migration^[15]^. There is evidence that changes in calcium-dependent cell adhesion molecules, such as cadherin-1 (CDH1), are associated with tumors. Cadherin downregulation can be used in the diagnosis and prognosis of epithelial cancer^[16]^. Cadherin-13 (CDH13) plays a regulatory role in tumor growth and promotes angiogenesis^[16, 17]^. In addition, tumor cells secrete high levels of angiogenic factors, which contribute to the generation of abnormal vascular networks, making inhibition of angiogenesis a key target for cancer therapy^[18]^.

In the cellular component category (Figure 5(b)), the majority of these differentially expressed proteins were secreted. In the molecular function category (Figure 5(c)), growth factor binding was overrepresented.

#### 3.2.4 Differences in protein expression between the complete resection group and the healthy control group at Day 30

Compared to the healthy control group, there were 8 differential proteins (shown in Table 4) in the complete resection group on Day 30 after surgery, of which 3 proteins were upregulated and 5 proteins were downregulated. These proteins are not sufficient to enrich biological processes. Random grouping statistical analysis showed that an average of approximately 3.77 differential proteins were identified out of 125 random grouping results, indicating a false-positive rate of 47.125% for differential proteins. This suggests that the physiological condition of the complete resection group was the less different from that of the healthy control group after 30 days of recovery.

**Table 4.** Differential proteins on Day 30 between the complete resection group and the healthy control group

| Protein Accessions | Protein Descriptions | Human ortholog | *p* value | Fold change |
| --- | --- | --- | --- | --- |
| Q922U2 | Keratin, type II cytoskeletal 5 | P13647 | 4.10E-02 | 2.102 |
| P60710 | Actin, cytoplasmic 1 | P60709 | 1.63E-02 | 1.695 |
| O55186 | CD59A glycoprotein | P13987 | 4.71E-02 | 1.502 |
| P11276 | Fibronectin | P02751 | 1.79E-02 | 0.617 |
| P0CW02 | Lymphocyte antigen 6C1 | - | 2.46E-02 | 0.609 |
| P06869 | Urokinase-type plasminogen activator | P00749 | 2.43E-02 | 0.576 |
| Q05793 | Basement membrane-specific heparan sulfate proteoglycan core protein | P98160 | 1.04E-02 | 0.526 |
| Q91WR8 | Glutathione peroxidase 6 | P59796 | 8.99E-03 | 0.480 |

## 3 Conclusion

Our results revealed that urine proteomics could distinguish between tumor-bearing mice and tumor-resected mice. These findings may provide a new strategy for clinical studies.

## Notes

### Competing Interest Statement

The authors have declared no competing interest.

